# Modeling the SARS-CoV-2 mutation based on geographical regions and time

**DOI:** 10.1101/2021.08.11.455941

**Authors:** Bomin Wei, Yuzhe Sun, Xiang Gong

## Abstract

The Coronavirus Disease 2019 (COVID-19) epidemic was first detected in late-December 2019. So far, it has caused 203,815,431 confirmed cases and 4,310,623 deaths in the world. We collected sequences from 150,659 COVID-19 patients. Based on the previous phylogenomic analysis, we found three major branches of the virus RNA genomic mutation located in Asia, America, and Europe which is consistent with other studies. We selected sites with high mutation frequencies from Asia, America, and Europe. There are only 13 common mutation sites in these three regions. It infers that the viral mutations are highly dependent on their location and different locations have specific mutations. Most mutations can lead to amino acid substitutions, which occurred in 3/5’UTR, S/N/M protein, and ORF1ab/3a/8/10. Thus, the mutations may affect the pathogenesis of the virus. In addition, we applied an ARIMA model to predict the short-term frequency change of these top mutation sites during the spread of the disease. We tested a variety of settings of the ARIMA model to optimize the prediction effect of three patterns. This model can provide good help for predicting short-term mutation frequency changes.

## INTRODUCTION

The Coronavirus Disease 2019 (COVID-19) has become a severe epidemic, claiming 203,815,431 cases and 4,310,623 deaths worldwide until August 2021^1^. Modern transportation and more frequent personnel exchanges have accelerated the spread of COVID-19. The second outbreak of this disease has plagued many countries where the epidemic is not serious. One of the main reasons is the long-distance migration of the mutated viral host, causing the new types of COVID-19 to spread across regions.

The COVID-19 is caused by a novel evolutionary divergent RNA virus, called severe acute respiratory syndrome coronavirus 2 (SARS-CoV-2), which triggers a respiratory tract infection and spreads mainly through person-to-person contact^2^. The genetic information of SARS-CoV-2 mutates much more dramatically than DNA due to RNA viruses’ mechanisms^3^. Until now, the new mutations of the virus. The worldwide outbreak happens to provide good environments for SARS-CoV-2 mutations. The accumulation of these mutations may cause the COVID-19 to develop in an uncertain direction, which will have a huge impact on society and personal life ^4^.

According to other epidemiological studies, mutations in the genome of an epidemic will be inherited from the spreader to the next generation of patients. The spread of the disease generally has regional characteristics, which leads to the diversity of the genome with regional traits. Thus, the SARS-CoV-2 genome mutation should have divergent mutation patterns in different geographic locations. Our purpose is to study the mutation patterns of SARS-CoV-2 in the world and try to predict the trend of the mutation so that to provide a reference for other researchers and may be helpful to the choice of the vaccine. To compare the mutation patterns quantitatively, we consider the cumulative mutation frequency on every single genome site as a time series. Previous studies in the aspect of mutation patterns and mutation predictions had focused on the protein mutation and functional changes of a certain genetic variant^3^, and there is no overview on the whole genomic sequence. Also, the studies had only limited prediction powers. They had not predicted all unknown mutations which could escape the vaccines already developed. Therefore, in this work we address this problem by employing a robust time series model, known as the ARIMA model for genetic mutation predictions. Finally, we shall develop a visualizing system to model the SARS-CoV-2 mutation trend based on geographical regions and time.

## METHODS

### Sample data filtering

As of January 31, 2021, the China National Center for Bioinformation (CNCB) database hosted 528,611 SARS-COV-2 sequences. Low-quality data (as assessed by the database) was removed. The data that required authorization by the submitting agencies were excluded. At last, 184,475 entries of the raw sequence were download^5^. An entry of raw sequence data directly downloaded from the CNCB database does not contain all the necessary metadata information such as host, sampling location, or date. The information was documented in a set of sequence metadata information, which needed to be downloaded separately. We performed a complete paring search to match the raw sequence data to the sequence metadata information and obtained 183,850 data entries with metadata information.

We discarded irrelevant metadata information and kept the following information for analysis and modeling: sequence name, detection time, detection region, base name before mutation, location of mutation site, base name after mutation. The base names and site also guarantee an easy retrieval of information needed for amino acid substitution analysis.

The data was divided based on the continent of the detection region and then sorted by time. We performed a further data cleaning and dropped those data with the non-standard format or incomplete metadata information. For example, we dropped a group of data reported from Japan which labeled detection time with only the year and the month but not the date. After the cleaning, 150,659 data entries remained.

We focused on data from Europe, Asia, and North America due to sufficient data quantity and high data quality in these regions. For analysis and modeling, we selected those sites which have more than 0.1% mutation rate on the last day and larger than 10% mutation rate on average.

### Development of prediction model

For the prediction model, the ARIMA (i.e, Auto Regressive Integrated Moving Average) model is used because of its advantages in time series forecasting. An ARIMA model contains three parameters – *p, d*, and *q*, or written as ARIMA(*p,d,q*). The *p* represents the number of lag observations in the auto-regression (AR) part of the model, indicating a relation between an observation (or data) to the past observations. The *q* represents the size of the moving average window in the moving average (MA) part of the model, indicating the relation between an observation to the past error. The *d* represents the integration order of the I(Integrated) part of the model, indicating the number of times that the raw observations are differenced. If we let *ŷ* be the *d*^*th*^ difference of Y (the observation or data), then the model can be expressed as *ŷ*_*t*_ = *μ*+*ϕ*_1_*y*_*t*−1_+…+*ϕ*_*p*_*y*_*t*−*p*_ − *θ*_1_*e*_*t*−1_ −… − *θ*_*q*_*e*_*t*−*q*_.

We developed an automatic parameter scan for 87 groups of data so that each mutation site had its model parameters fitted independently. For the mutation rate data of each site, the program tests 5 different parameters for *p* and *q*, respectively, and number 1 or 2 for parameter *d*. In total, there were 50 parameter combinations tested for every of the high-frequency sites. Values of *p* are first determined from the ACF (autocorrelation function) of the mutation rate. The best parameter set (*p, q, d*) which has the lowest MSE value is used as parameter for the prediction.

## RESULTS

### Data collection and description

Since COVID-19 has been circulating for more than a year, virus samples from many countries have been sequenced. The samples range from dozens of countries, and the sampling time covers several months. This gives us great convenience to study gene mutations in different regions and their changing trends over time. We downloaded mutation data of the disease from a public database: China National Center for Bioinformation, 2019nCoVR (https://bigd.big.ac.cn/ncov/?lang=en)^5^. In total, mutation information from 150,659 patients was downloaded from the database. We used the SARS-CoV-2 sequence, NC_045512, of the first COVID-19 patient as the reference sequence in this study. The genome is with the length of 29,903 bp ss-RNA ^6^. We obtained the corresponding sample location (country) and sampling time as well. According to the sample source, 74% of the samples are from Europe, 15% of the samples are from North America, and 7% of the samples are from Asia (Figure 1A). In terms of time span, the sample starts from January 2020 to January 2021 (Figure 1B). The samples in Europe increase dramatically after July 2020, while samples in Asia and North America slightly increase from April 2020. Samples from Europe accounted for the largest proportion. Although some countries, such as the USA have carried out large-scale COVID-19 positive tests for their citizens, only a few whole-genome sequences were available. Although the proportion of the sequences in America and Asia are low, there are still 22,599 sequences in America and 10,546 sequences in Asia, and the mutation frequency calculation should be precise.

**Figure 1.**
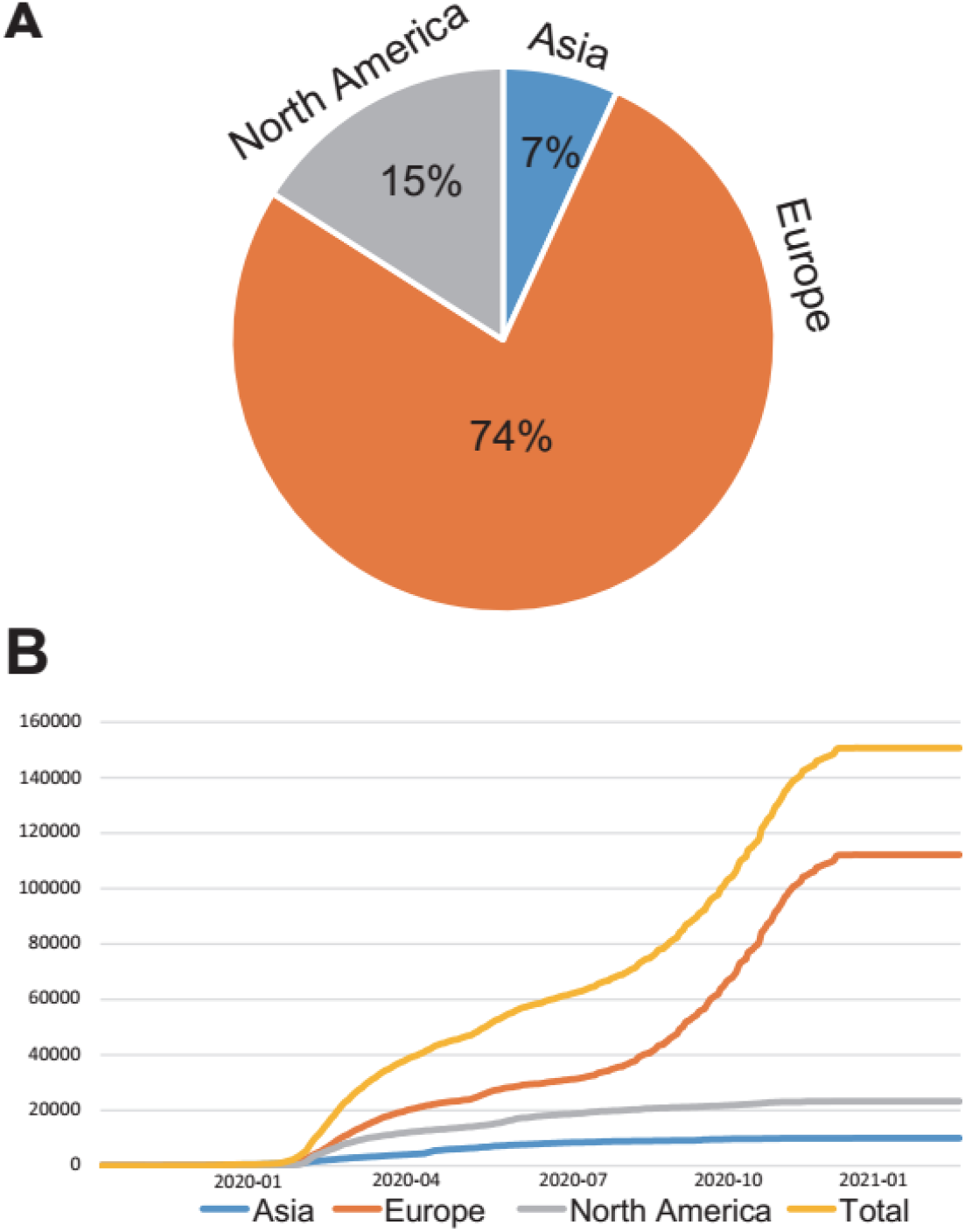
Distribution of downloaded SARA-Cov-2 sequence mutation data.

### Different patterns of single nucleotide polymorphisms (SNPs) in Asia, Europe, and North America

We focused on the single-site mutation (SNP) and calculated the frequencies of all mutation sites (Figure 2). The mutations are ubiquitous in most regions of the genome and the sites with the highest mutation frequencies are in Polyprotein (ORF1ab), S protein, ORF3a, M protein, ORF8, N protein, ORF10 (Figure 2).

**Figure 2.**
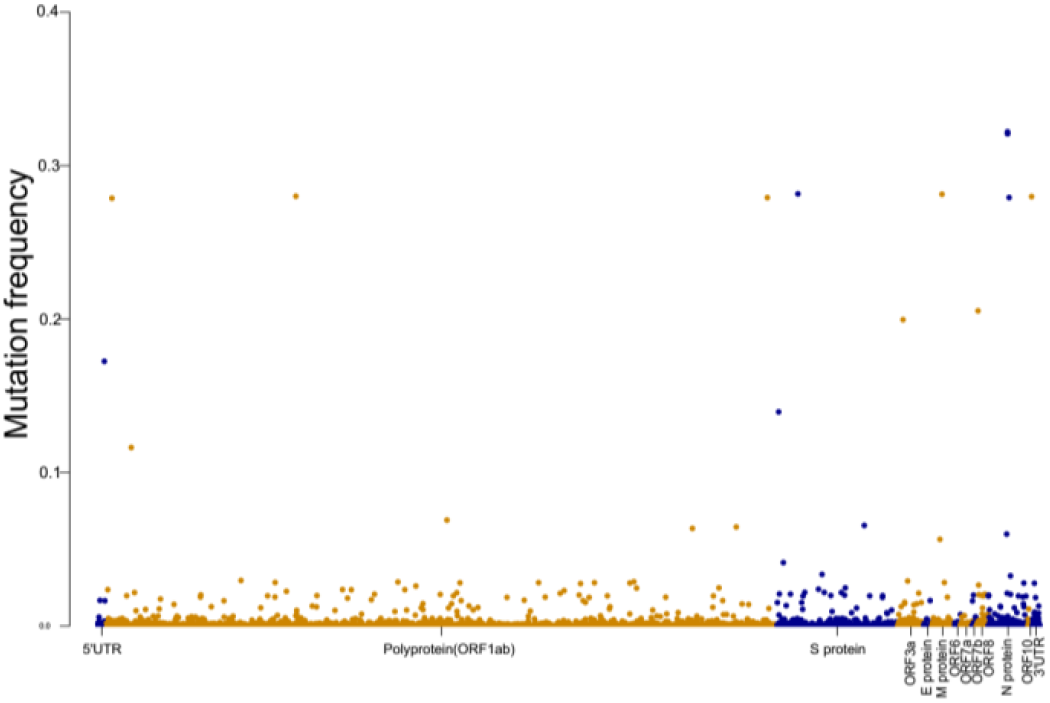
Distribution of SNPs on SARS-CoV-2 genome.

We selected the sites with top mutation frequencies in Asia, Europe, and North America. There are 27 sites in Asia samples, 30 sites in Europe samples, and 30 sites in North America samples (Figure S1). Only 13 sites are shared by all three regions suggesting that the mutation patterns may be different in three regions. We also calculated the mutation frequency changes with time (Figure S2). Three main clusters can be observed: 1) The mutation frequency gradually increases with time, and finally reaches a plateau. This pattern likes a classic logistic curve. 2) The mutation frequency has a high peak at the beginning of the outbreak, and then the frequency decreases to a low level with time. 3) The mutation frequency has a high peak at the beginning of the outbreak, and then the frequency decreases with time but maintains a high level. Cluster 1 takes for a little more than 30% of each group. Cluster 2 takes about 30% of Asia and Europe samples, while it takes more than two-thirds of North America samples. Cluster 3 takes around 30% in Asia and 20% in Europe (Figure S1). It confirms the different patterns in Asia, Europe, and North America samples.

### Most of the mutation can lead to amino acid substitution

Non-synonymous and synonymous mutations can lead to different results. Thirty-eight SNPs in the SARS-CoV-2 genome can contribute to amino acid substitution (Table S1). Three genes, ORF1ab, Spike protein, and N protein, contain the most non-synonymous mutations (Table 1). Spike protein mediates host cell receptor recognition and binding. It is the key for vaccine design and development against SARS-CoV-2 infection^7^. The enrichment of SNPs in the Spike protein gene could lead to the subsequent evolution of the virus. Three regions display different mutation directions which makes the prediction of the mutation valuable (Table 1).

**Table 1.** Non-synonymous mutations enriched in three gene loci.

| Gene | Asia | Europe | America |
| --- | --- | --- | --- |
| ORF1ab | 9 | 9 | 11 |
| Spike protein | 1 | 3 | 2 |
| N protein | 4 | 4 | 4 |
| Cluster (1/2/3) | 7/2/5 | 6/5/5 | 7/10/0 |

### Model of predicting the mutation frequency

Based on a regression method, we developed a model to predict the frequency change with time in a short period (Figure S1). The MSE values represent the accuracy and stability of the model (Table 2). The model proposes the mutation trends of each site in three regions. The method considers multiple factors: disease outbreak region, number of days in the training set, and number of days for out-of-sample prediction, and predicted the mutation frequency for specific sites. It could be a reference for researchers on different continents.

**Table 2.** MSE of the prediction model.

| MSE | Average ($-\log_{10}$) | SD |
| --- | --- | --- |
| Asia | 6.97 | 0.95 |
| Europe | 4.62 | 1.50 |
| America | 6.93 | 0.86 |

The mutations in Spike protein may affect the off-target effect of the vaccine. We selected a few genome positions for detailed modeling analysis in the range 21,000 to 26,000, which is believed to encode the spike protein.

We selected the A23403G mutation which is the D614G spike protein mutation. We used the ARIMA model to predict the short-term mutation frequency. The training data set was defined from the first day of non-zero mutation rate to the 200^th^ day. We tested three ARIMA model parameters (*p, d, q*), and d = 2 followed the rising trend of the real mutation rate data (Figure S3). The ARIMA model parameters (*p, d, q*) = (2, 2, 3) were fitted on the training set and the optimized parameters were selected to minimize the MSE on the test set from the 201^st^ day to the 280^th^ day, between the out-of-sample forecast and the real data (Figure 3). When the forecast is performed in a much longer period from the 201^st^ day to the 360^th^ day, the ARIMA model can only capture the rising trend but failed to fit the plateau of the real data (Figure S4). Another genome position is 22,227 (C-to-T, also called A222V), of which the mutation rate shows a logistic curve and reaches a steady rate beyond the 160^th^ day. ARIMA models with parameters *d* = 1 can capture this steady rate and the best forecast occurs at (*p, d, q*) = (3, 1, 1) (Figure 4). The training set of this position had a different length compared to position 23403 because the mutation rate stabilizes much faster, and we used the time range from the first date of non-zero mutation rate to the 130^th^ day after the first day. The optimized parameters are sensitive to the choice of the training set length, which invites further study into the modeling stability.

**Figure 3.**
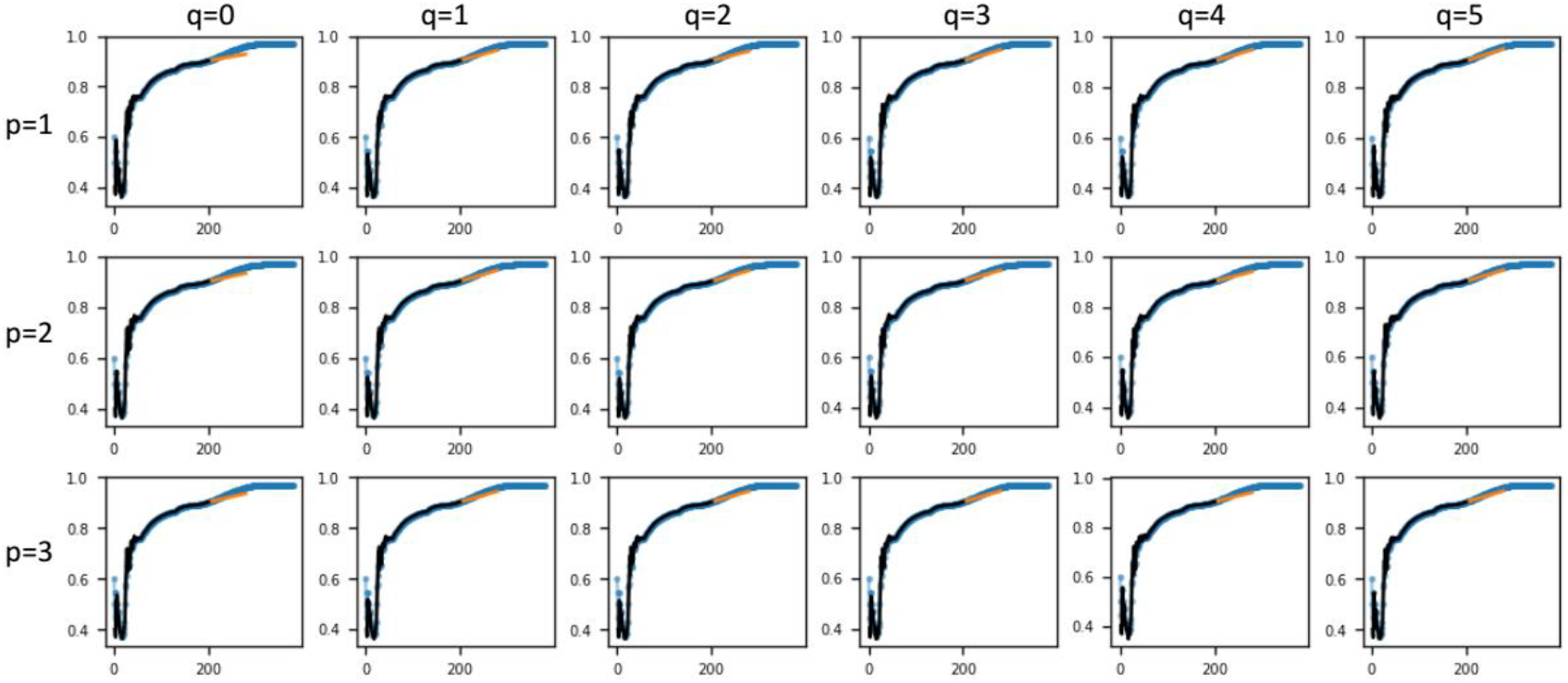
The ARIMA model fitting and forecast of the genome position 23403. The matrix of plots (*p, q*) all have a fixed difference order *d*=2. The real data (blue curve) and the in-sample prediction (black curve) for the training data set (from the date of the first mutation report to the 200th day after that data) fits well, and the out-of-sample forecast from the 201st to the 280th day (orange curve) achieves minimum MSE with parameters (*p, d, q*) = (2, 2, 3).

**Figure 4.**
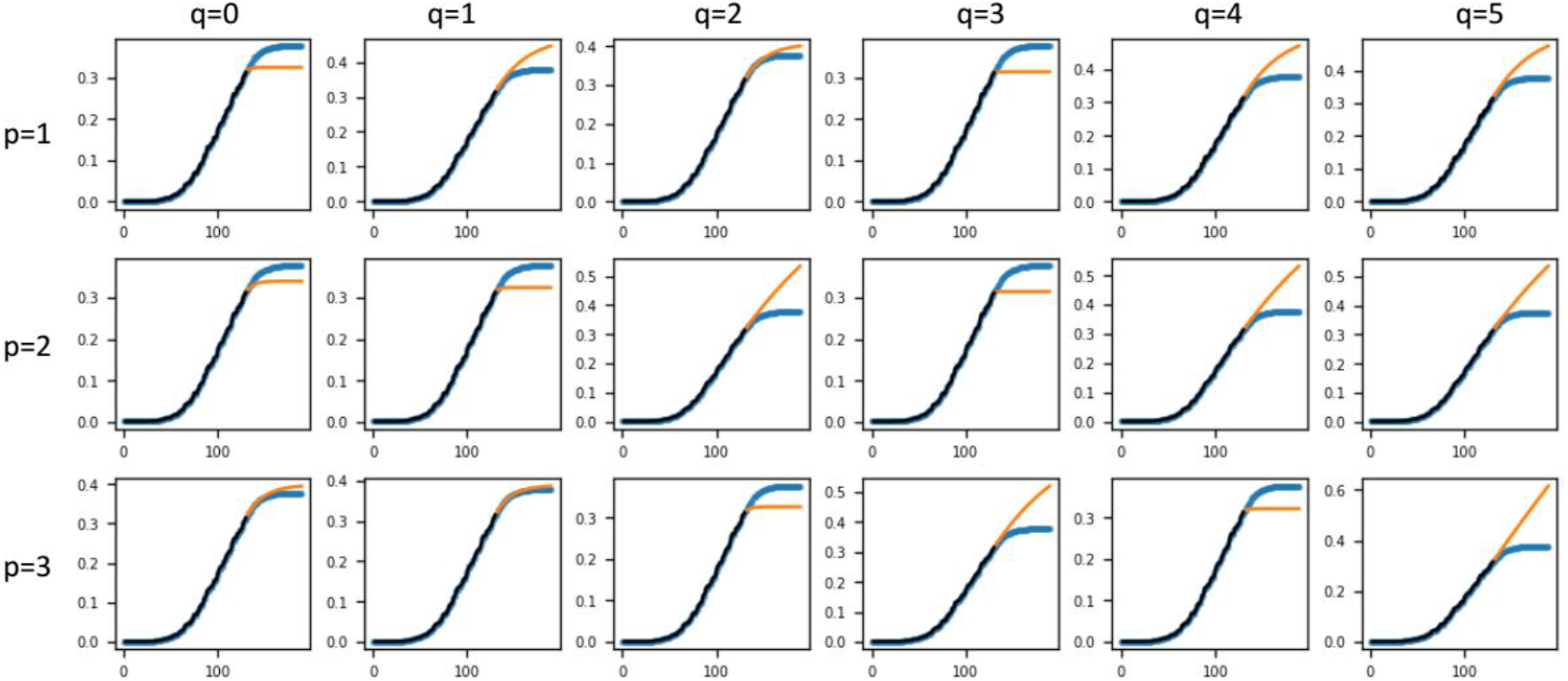
ARIMA model fitting and forecast of the genome position 22227. The matrix of plots (*p,q*) all have a fixed difference order *d*=1. The real data (blue curve) and the in-sample prediction (black curve) for the training data set fits well, and the short period forecast (orange curve) from 131^st^ day to the 210^th^ day reproduces the correct real data for (*p, d, q*) = (3, 1, 1).

Based on the similarity of genomes and the prediction of mutation trends, we hope that our work can provide an alternative reference for residents to choose vaccines produced in different brands and different countries. Overall, we expect our approach to be a fundamental solution in the literature and contribute as reliable quantitative benchmarks.

## Discussion

We focused on the Spike protein which is the most important region in the SARS-CoV-2 genome related to human immune response (Figure 5) ^8^. The angiotensin-converting enzyme 2 (ACE2) proteins can bind the glycosylated S proteins on the surface of SARS-CoV-2 and mediates the viral cell entry^9^. The SARS-CoV-2 S protein consists of 1273 amino acids, including a signal peptide in the N-terminus, the S1 and S2 subunits (Figure 5). In the S1 subunit, a receptor-binding domain (RBD) is responsible for binding the ACE2. Fusion peptide (FP) plays an essential role in mediating membrane fusion^10^. HR1 and HR2 form the six-helical bundle (6-HB) with the entry function of the S2 subunit^11^. The transmembrane (TM) domain of the S protein anchors in the viral membrane.

**Figure 5.**
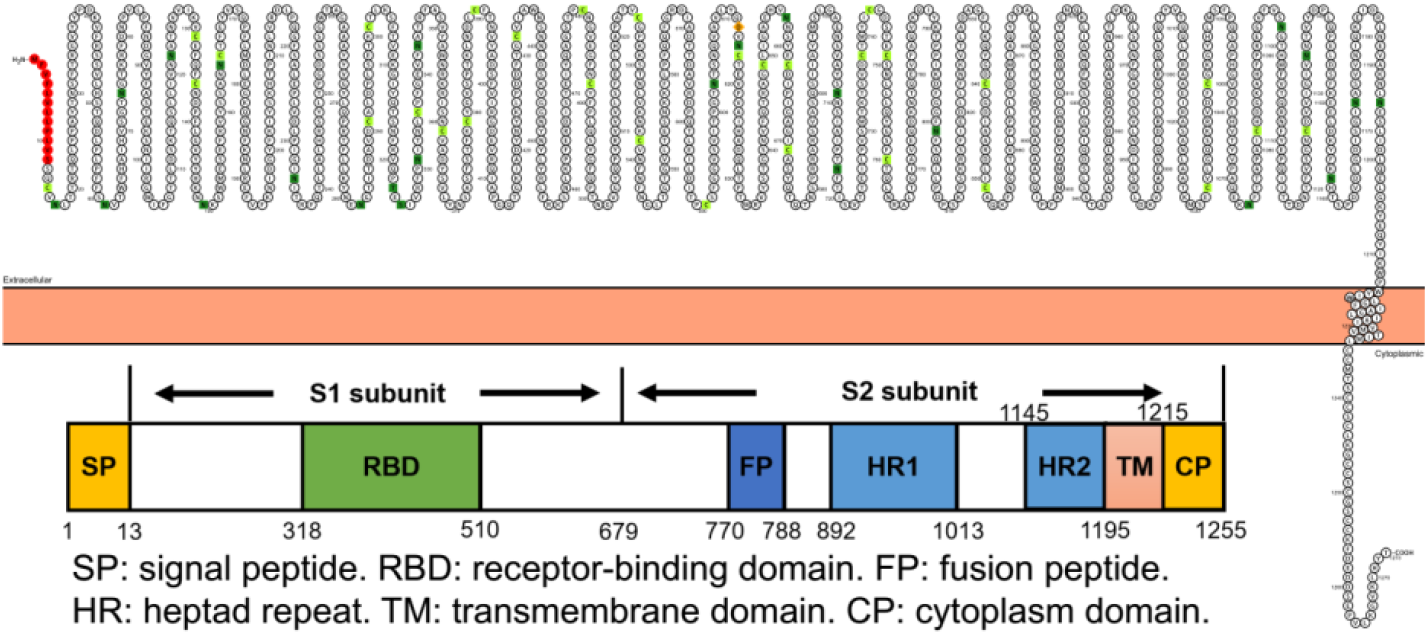
Structure of Spike protein in SARS-CoV-2 genome.

We found three patterns of the SARS-Cov-2 mutation patterns. We have tested the ARIMA model in cluster 1 and the model exhibits good performance in the short-term prediction. We further applied the ARIMA model mutation sites of the other two clusters. The genome mutation C27046T (T175M) is a typical cluster 2 cases in Europe. The prediction for the training data set fits well with the real data, and the short-term prediction with (*p, d, q*) = (3, 1, 1) in the range from 116^th^ day to the 176^th^ day reproduces the correct real data (Figure S5). Another typical cluster 3 case in Asia is C28311T (P13L). With the settings (p, d, q) = (3, 1, 2), the short-term prediction from the 200^th^ day to the 260^th^ day reproduces the correct real data (Figure S6).

In summary, we collected SARS-Cov-2 sequence mutations and summarized 3 mutation patterns. We screened out non-synonymous mutation sites and found that the patterns of mutations on different continents are different. We fitted the mutation frequencies to the ARIMA model, and the model can forecast well in the short term. It would be a good basis for other studies of COVID-19 in terms of genomic mutation patterns and prediction.

---

**Figure S1.**
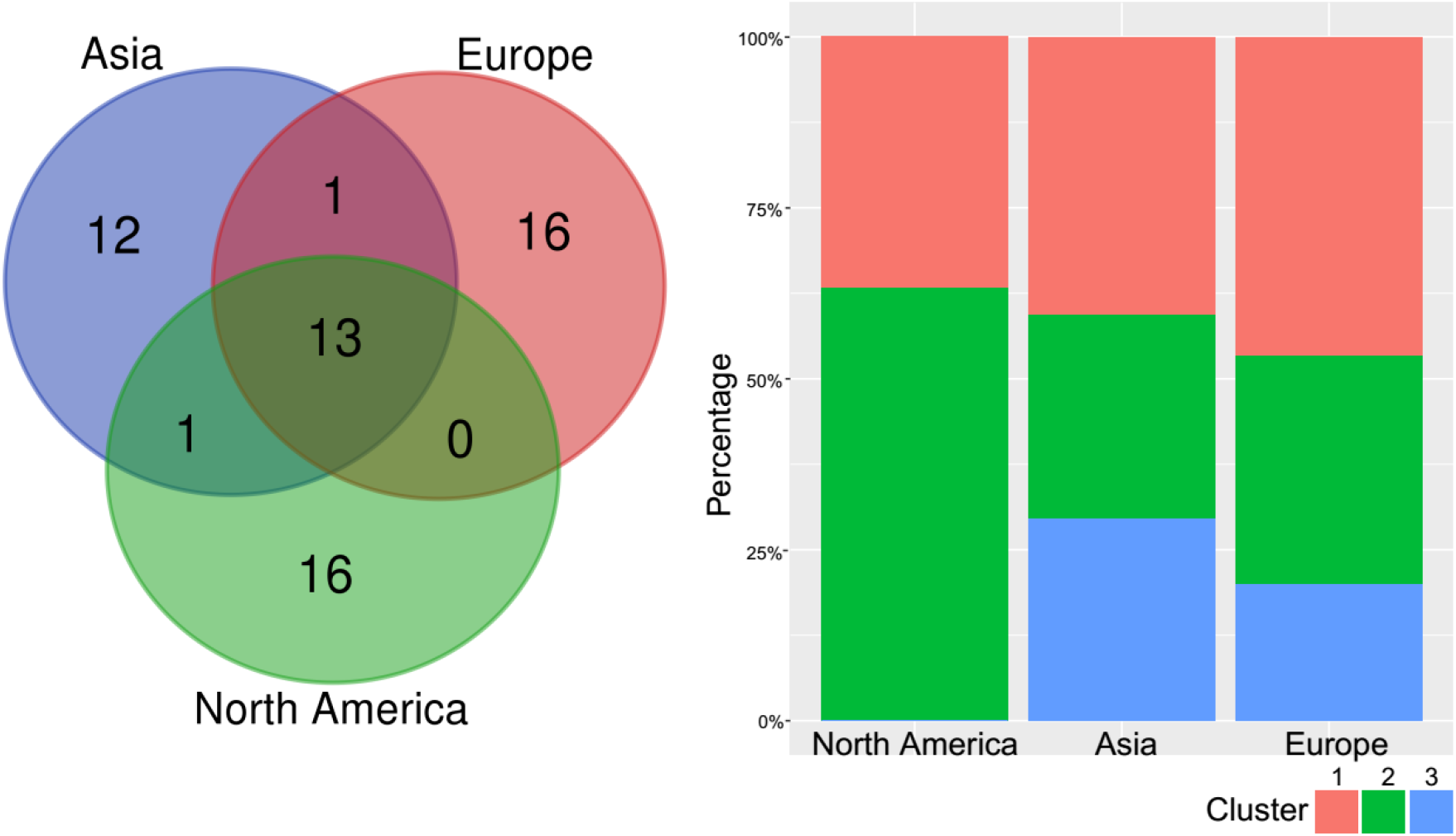
Sites with top mutation frequencies in Asia, Europe, and North America.

**Figure S2.**
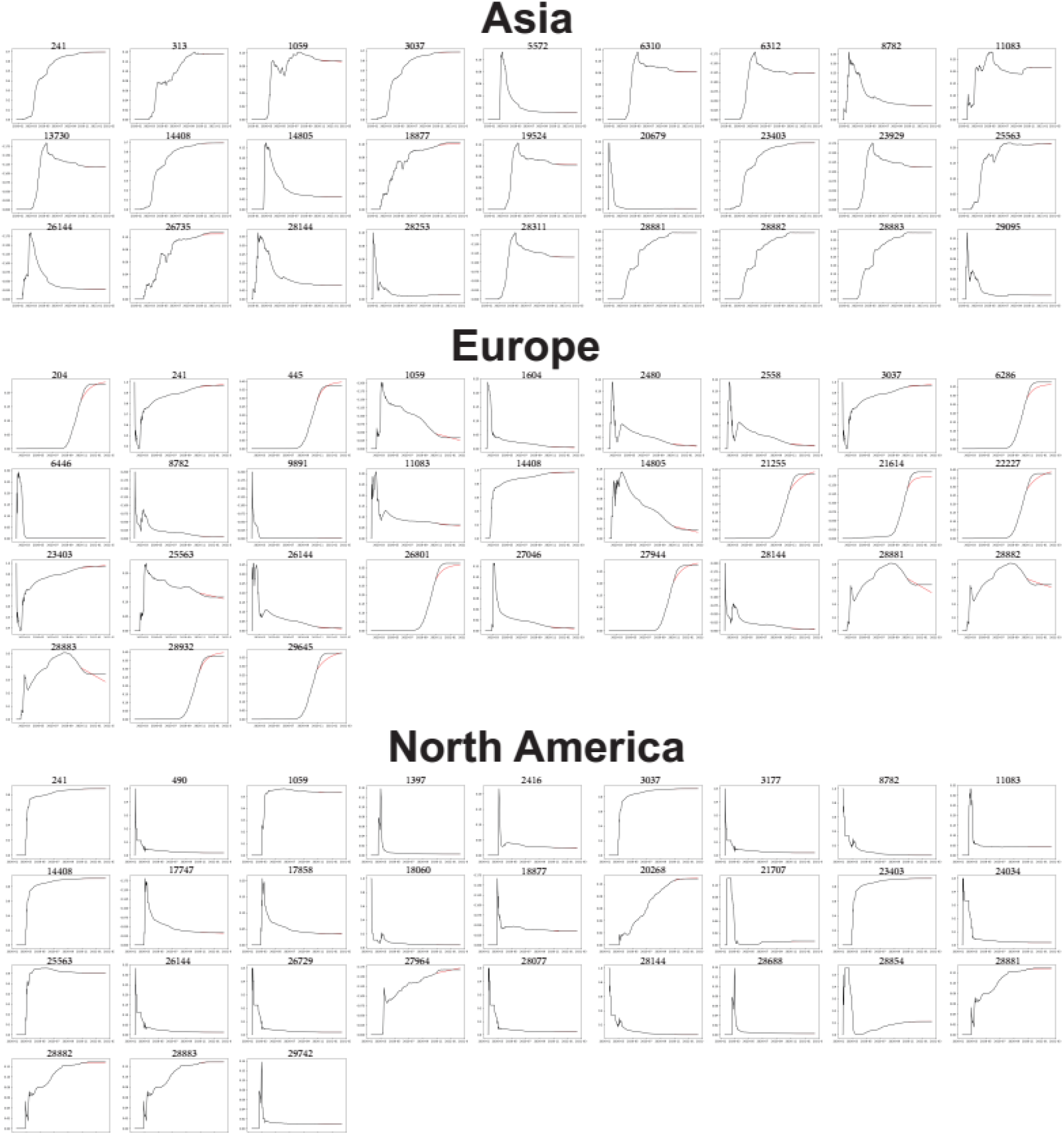
Frequencies and prediction of top mutated sites in Asia, Europe, and North America.

**Figure S3.**
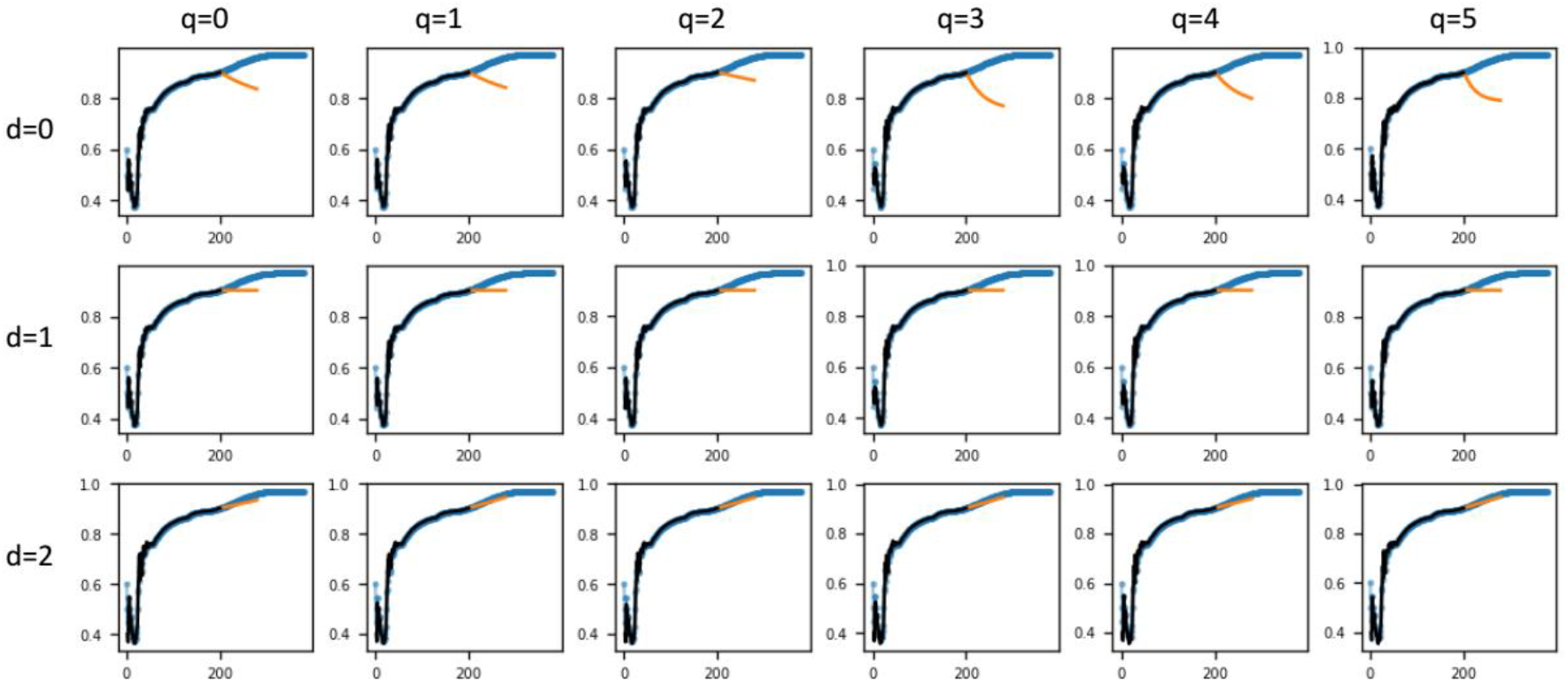
ARIMA model fitting and forecast of the genome position 23403. The matrix of plots (d, q) all have a fixed p=2. The real data (blue curve) and the in-sample prediction (black curve) for the training data set fits well, and the out-of-sample forecast from the 201^st^ to the 280^th^ day (orange curve) only follow the correct trend of real data at d=2.

**Figure S4.**
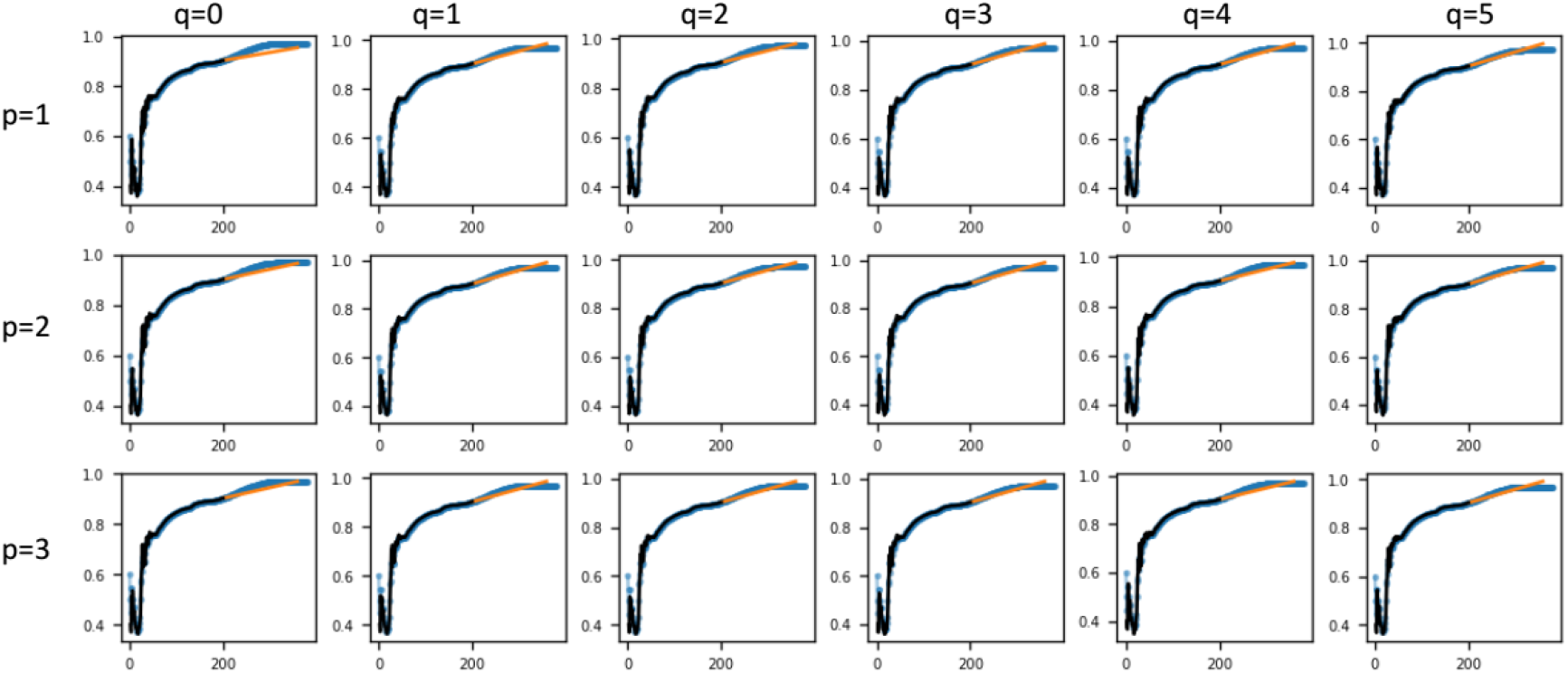
ARIMA model fitting and forecast of the genome position 23403. The matrix of plots (p, q) all have a fixed difference order d=2. The real data (blue curve) and the in-sample prediction (black curve) for the training data set fits well, but the long period forecast (orange curve) from 201^st^ day to the 360^th^ day shows that none of the parameter combinations of the ARIMA model can reproduce the plateau beyond 300^th^ day after the mutation onset.

**Table S1.** Most mutated sites in the SARS-CoV-2 genome.

| Position | Gene | SNP | Codon change | Amino acid position | Amino acid change | Non-synonymous |
| --- | --- | --- | --- | --- | --- | --- |
| 313 | ORF1ab | C -> T | CTC->CTT | 16 | L->L | No |
| 445 | ORF1ab | T -> C | GTT->GTC | 60 | V->V | No |
| 490 | ORF1ab | T -> A | GAT->GAA | 75 | D->E | yes |
| 1059 | ORF1ab | C -> T | ACC->ATC | 265 | T->I | yes |
| 1397 | ORF1ab | G -> A | GTA->ATA | 378 | V->I | yes |
| 2416 | ORF1ab | C -> T | TAC->TAT | 717 | Y->Y | No |
| 2480 | ORF1ab | A -> G | ATT->GTT | 739 | I->V | yes |
| 2558 | ORF1ab | C -> T | CCA->TCA | 765 | P->S | yes |
| 3037 | ORF1ab | C -> T | TTC->TTT | 924 | F->F | No |
| 3177 | ORF1ab | C -> T | CCT->CTT | 971 | P->L | yes |
| 5572 | ORF1ab | G -> T | ATG->ATT | 1769 | M->I | yes |
| 6286 | ORF1ab | C -> T | ACC->ACT | 2007 | T->T | No |
| 6310 | ORF1ab | C -> A | AGC->AGA | 2015 | S->R | yes |
| 6312 | ORF1ab | C -> A | ACA->AAA | 2016 | T->K | yes |
| 6446 | ORF1ab | G -> T | GTT->TTT | 2061 | V->F | yes |
| 8782 | ORF1ab | C -> T | AGC->AGT | 2839 | S->S | No |
| 9891 | ORF1ab | C -> T | GCT->GTT | 3209 | A->V | yes |
| 11083 | ORF1ab | G -> T | TTG->TTT | 3606 | L->F | yes |
| 13730 | ORF1ab | C -> T | CTA->TTA | 4489 | A->L | yes |
| 14408 | ORF1ab | C -> T | CTA->TTA | 4715 | P->L | yes |
| 14805 | ORF1ab | C -> T | ACT->ATT | 4847 | Y->I | yes |
| 17747 | ORF1ab | C -> T | CTG->TTG | 5828 | P->L | yes |
| 17858 | ORF1ab | A -> G | ATG->GTG | 5865 | Y->V | yes |
| 18060 | ORF1ab | C -> T | TCT->TTT | 5932 | L->F | yes |
| 18877 | ORF1ab | C -> T | GTC->GTT | 6204 | C->V | yes |
| 19524 | ORF1ab | C -> T | TCG->TTG | 6420 | L->L | No |
| 20268 | ORF1ab | A -> G | TAG->TGG | 6668 | L->W | yes |
| 21255 | ORF1ab | G -> C | CGT->CCT | 6997 | A->P | yes |
| 21614 | Spike protein | C -> T | CTT->TTT | 18 | L->F | yes |
| 21707 | Spike protein | C -> T | CAT->TAT | 49 | H->Y | yes |
| 22227 | Spike protein | C -> T | GCT->GTT | 222 | A->V | yes |
| 23403 | Spike protein | A -> G | GAT->GGT | 614 | D->G | yes |
| 23929 | Spike protein | C -> T | TAC->TAT | 789 | Y->Y | No |
| 24034 | Spike protein | C -> T | AAC->AAT | 824 | N->N | No |
| 25563 | ORF3a | G -> T | CAG->CAT | 57 | Q->H | yes |
| 26144 | ORF3a | G -> T | GGT->GTT | 251 | G->V | yes |
| 26729 | M protein | T -> C | GCT->GCC | 69 | A->A | No |
| 26735 | M protein | C -> T | TAC->TAT | 71 | Y->Y | No |
| 26801 | M protein | C -> G | CTC->CTG | 93 | L->L | No |
| 27046 | M protein | C -> T | ACG->ATG | 175 | T->M | yes |
| 27944 | ORF8 | C -> T | CAC->CAT | 17 | H->H | No |
| 27964 | ORF8 | C -> T | TCA->TTA | 24 | S->L | yes |
| 28077 | ORF8 | G -> C | GTG->CTG | 62 | V->L | yes |
| 28144 | ORF8 | T -> C | TTA->TCA | 84 | L->S | yes |
| 28253 | ORF8 | C -> T | TTC->TTT | 120 | F->F | no |
| 28311 | N protein | C -> T | CCC->CTC | 13 | P->L | yes |
| 28688 | N protein | T -> C | TTG->CTG | 139 | L->L | no |
| 28854 | N protein | C -> T | TCA->TTA | 194 | S->L | yes |
| 28881 | N protein | G -> A | AGG->AAA | 203 | R->K | yes |
| 28882 | N protein | G -> A | AGG->AAA | 203 | R->K | yes |
| 28883 | N protein | G -> C | GGA->CGA | 204 | G->R | yes |
| 28932 | N protein | C -> T | GCT->GTT | 220 | A->V | yes |
| 29095 | N protein | C -> T | TTC->TTT | 274 | F->F | no |
| 29645 | ORF10 | G -> T | GTA->TTA | 30 | V->L | yes |

**Figure S5.**
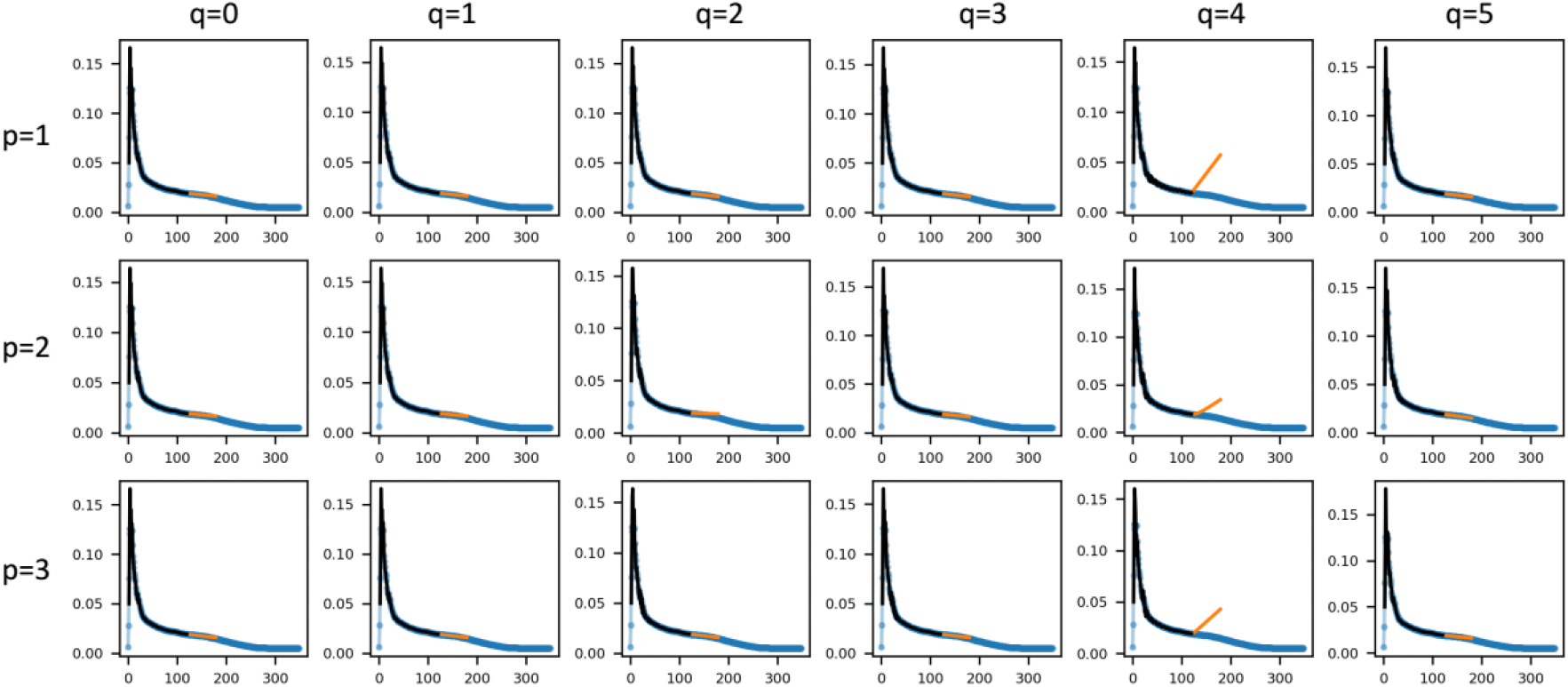
ARIMA model fitting and forecast of the genome position 27046, a typical cluster 2 case in Europe. The matrix of plots (p, q) all have a fixed difference order d=1. The real data (blue curve) and the out-of-sample prediction (orange curve) for the training data set fits well, and the short period forecast (orange curve) from 116^th^ day to the 176^th^ day reproduces the correct real data for (p, d, q) = (3, 1, 1).

**Figure S6.**
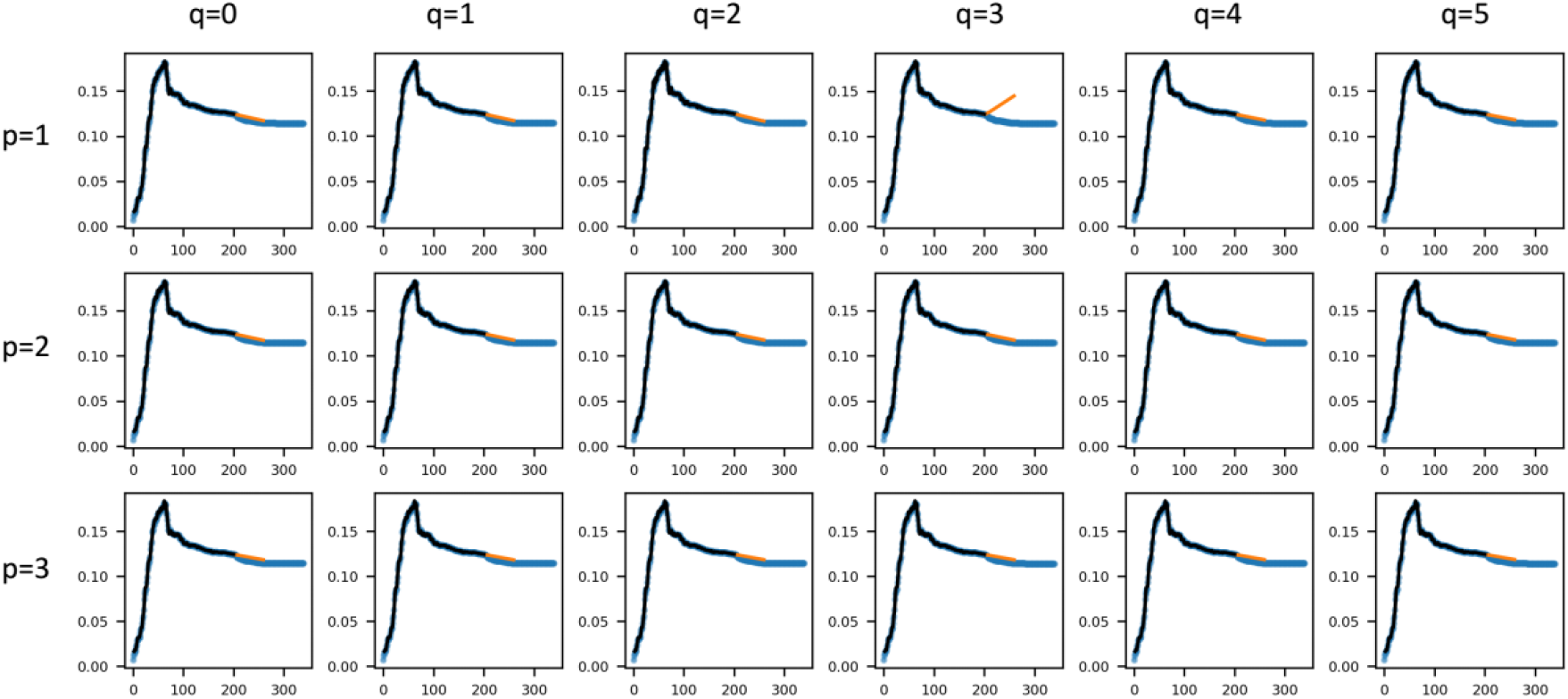
ARIMA model fitting and forecast of the genome position 28311, a typical cluster 3 case in Asia. The matrix of plots (p, q) all have a fixed difference order d=1. The real data (blue curve) and the out-of-sample prediction (orange curve) for the training data set fits well, and the short period forecast (orange curve) from 200^th^ day to the 260^th^ day reproduces the correct real data for (p, d, q) = (3, 1, 2).

## Notes

### Competing Interest Statement

The authors have declared no competing interest.

